# Wide-Angle, Monocular Head Tracking using Passive Markers

**DOI:** 10.1101/2021.09.01.458583

**Authors:** Balazs P. Vagvolgyi, Ravikrishnan P. Jayakumar, Manu S. Madhav, James J. Knierim, Noah J. Cowan

**Author notes:** Corresponding author: Email address (Balazs P. Vagvolgyi).

## Abstract

Camera images can encode large amounts of visual information of an animal and its environment, enabling high fidelity 3D reconstruction of the animal and its environment using computer vision methods. Most systems, both markerless (e.g. deep learning based) and marker-based, require multiple cameras to track features across multiple points of view to enable such 3D reconstruction. However, such systems can be expensive and are challenging to set up in small animal research apparatuses.

We present an open-source, marker-based system for tracking the head of a rodent for behavioral research that requires only a single camera with a potentially wide field of view. The system features a lightweight visual target and computer vision algorithms that together enable high-accuracy tracking of the six-degree-of-freedom position and orientation of the animal’s head. The system, which only requires a single camera positioned above the behavioral arena, robustly reconstructs the pose over a wide range of head angles (360^◦^ in yaw, and approximately ±120^◦^ in roll and pitch).

Experiments with live animals demonstrate that the system can reliably identify rat head position and orientation. Evaluations using a commercial optical tracker device show that the system achieves accuracy that rivals commercial multi-camera systems.

Our solution significantly improves upon existing monocular marker-based tracking methods, both in accuracy and in allowable range of motion.

The proposed system enables the study of complex behaviors by providing robust, fine-scale measurements of rodent head motions in a wide range of orientations.

## 1. Introduction

Organismal biology often relies on measurement and characterization of an animal’s anatomy, physiology, and behavior in the context of its external environment. Behavior represents the sum of motoric output of the animal, generated by a combination of internal neural dynamics and responses to external stimuli. Behavioral experimenters try to control and measure external stimuli presented to the animal. These measurements are typically performed at a rate and resolution appropriate to the time constants and scale of the corresponding biological variables. Movements on the scale of multiple body lengths can be measured using a variety of techniques. GPS [1, 2] and camera tracking [3, 4, 5, 6, 7, 8, 9, 10] are common approaches. Some tracking techniques are specific to peculiarities of individual species, for example acoustic triangulation for bats and whales and localization of electric fish using a grid of electrodes [11, 12, 13, 14, 15, 16, 17]. Fine-scale, sub-body-length movements are harder to track and sometimes require a more invasive approach in which markers or mechanical sensors are placed at key locations on the animals body.

Optical marker tracking has been a mainstay of behavioral research for decades. These systems typically use one or more cameras to capture images of the animal and its environment and hardware that implements computer vision algorithms to identify and track visual features on these images. Tracking can be performed offline or in real time. Offline trackers often employ sophisticated algorithms or less powerful processing hardware, since there are usually no major constraints on processing time. This approach is suitable for open-loop experiments, where tracking results do not affect the experiment. Real-time systems, however, usually involve a closed-loop experimental setup where tracking results are used to affect the animal or its environment in the same experiment. These tracking systems must complete their computations for each video frame prior to acquisition of the next frame, often requiring complex tracking algorithms or more expensive processing hardware.

### 1.1. Object Tracking Using Model-Based Approaches

Until recent years, most animal tracking methods relied on model-based computer vision approaches—a description of the appearance of image features to be identified (the models) and algorithms that search for features resembling these models on images. Model-based approaches are efficient and robust when the custom-built computational model fits the appearance of the corresponding image feature well; however they tend to fail when the model does not accommodate all possible feature appearances. It is usually easier to identify rigid visual features that look similar from every orientation. Conversely, soft or deformable features, or features that look different depending on point of view present a significant challenge for model-based systems.

Due to these limitations, model-based optical trackers often rely on simple markers that are easily identifiable instead of natural features of the environment or the animal. Commonly used markers include light emitting diodes (LEDs), retro-reflective spheres, high-contrast geometrical patterns and painted markers, all of which are detectable using simple image processing and their positions can be measured to high precision. Markers can in some cases be omitted if the model of the animal is simple enough (e.g. tracking position of a rodent on a bright background [18]), but these solutions tend to be less accurate.

The most commonly used markers, LEDs and retro-reflective spheres, represent a single point in space, and due to their rotational symmetry, they enable high accuracy position estimates from a wide range of view angles. However, resolving the orientation of a single, rotationally symmetric marker is impossible. Therefore, more complex visual targets, or multiple markers assembled in a known geometric configuration, are used when orientation tracking is required. Most widely used model-based optical trackers use a combination of easily identifiable image features to enable the calculation of orientation in addition to position. One commonly used system for animal tracking in laboratory settings is the multi-LED Video Tracker, where the relative position of two or three LEDs, shown in Fig. 1a, enables the calculation of the 2D orientation estimate.

**Figure 1:**
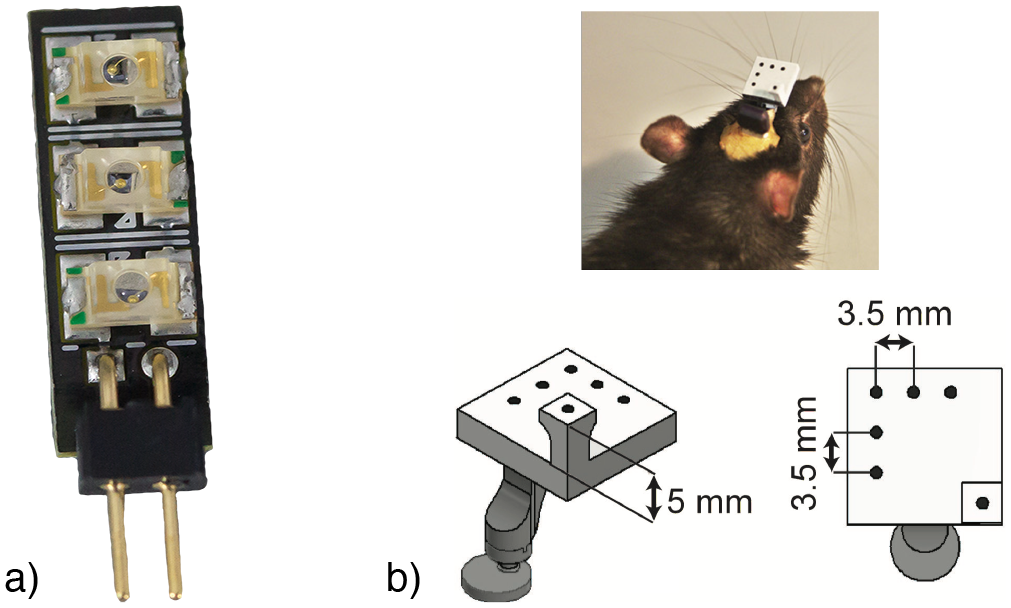
Commonly used marker systems: **(a)** 3-LED marker used for animal tracking in 2D. For rodents, the marker is often mounted on the back or the head of the animal. (Neuralynx Inc., Bozeman MT USA), **(b)** Marker used for 3D optical tracking of a rat’s head by Vanzella et al. [19]. The system tracks six black dots painted on a white plastic frame using a camera looking at the animal from above at a close distance.

Three-dimensional tracking can be achieved in two ways: using multiple cameras or by using a more complex marker design with enough information encoded in the appearance that 3D position and orientation (pose) can be resolved even from a single camera view. Using a single camera is experimentally simpler but the technique is more sensitive and complicated. Monocular pose estimation typically requires a complex marker with a known rigid geometry— such a marker is inherently larger than simple point markers. The marker size is dependent on the resolution of the camera; the resolution needs to be high enough so that computer vision algorithms are able to resolve individual features on the marker. Thus, higher camera resolutions allow smaller markers.

One popular class of monocular 3D optical trackers are ARTags (Augmented Reality Tags)—flat square shaped markers with a high-contrast pattern. The pattern in the middle identifies the marker and specifies its orientation out of the possible four configurations of a square. In this paper, for comparison tests, we use ArUco [20, 21] (Fig. 12), a widely used ARTag system in robotics and automation. While ARTag tracking is relatively robust and accurate, the plane of the marker must generally be facing the camera, and the range of viewing angles from which the ARTag can be detected is limited, since the visibility of the flat pattern diminishes when observed from a sharp angle. One common way to circumvent the view-angle limitation is to place multiple different ARTags on the sides of a rigid 3D object, for example multiple ARTags on each side of a cube [22, 23, 24], which enables the camera to see at least one marker at any orientation. Unfortunately, these complex markers featuring multiple ARTags are large and thus their use in animal behavioral tracking is limited.

Animal behavior researchers have developed other complex 3D marker designs for specific applications. In one system designed for mouse head tracking [19], the marker is a small 3D printed white plastic piece with six black dots arranged in a 3D pattern (Fig. 1b). While positioning one of the dots out of plane significantly improves 3D orientation estimation accuracy, this system still has similar disadvantage as ARTags: because the dots are all facing the same direction, it is not possible to detect them from behind the marker or from a sharp angle, severely limiting the range of usable orientations. Another system developed for tracking bats in flight [25] uses a set of LEDs in a custom arrangement that enables the reconstruction of the position and two of the three rotational degrees-of-freedom (DoF). While this system allows tracking in a wider range of angles, the configuration of LEDs still limits the visibility, and it is not suitable for applications in which all three rotational axes need to be resolved.

View-angle limitations are very common in optical tracking due to marker occlusions. Like ARTags, markers are often designed as one-sided, to prevent individual features on the marker from occluding one another. There exists a need for robust 3D tracking without these limitations, for example in tracking the head of a rodent. Rodent behaviors often involves a significant amount of head movement—scanning from side to side to evaluate the environment [26, 27, 28], making up and down motions to evaluate distance via parallax [29], and grooming where the head goes through a generally stereotypical set of movements as the animal cleans itself [30]. The most obvious solution to track these large ranges of head orientations is to use a multi-camera system. Commercially available multi-camera systems are expensive and often designed to measure motion on the scale of human kinematics. Cheaper, open-source implementations are often complicated to set up due to the need for camera extrinsic calibrations, camera sensor synchronization, increased computational requirements, and constraints in camera placement. In addition, installing multiple cameras with unobstructed views of the animal might restrict placement of other measurement and control apparatuses such as environment manipulation effectors, electrophysiological recording systems, or imaging equipment.

Avoiding a multi-camera solution requires a non-planar marker design that is small and detectable from most orientations, and a corresponding tracking algorithm that can handle the situation when only a subset of the marker’s features is visible due to occlusions. Faessler et al. [31] developed a marker-based tracker that uses 4–5 infrared (IR) LEDs mounted on the body of a flying drone in a configuration that enables at least four LEDs to be visible from an extended range of orientations. As the LEDs all look identical from the camera’s perspective and some of them may be occluded, the correspondence between the LEDs and the observed dots on the image need to be determined before performing 3D pose estimation using the Perspective-n-Point (PnP) algorithm. They use a brute force combinatorial method to find this correspondence. Their algorithm, in theory, is able to accommodate a more than five LEDs which would in increase the range of supported orientations; however the exponentially increasing combinatorial complexity of the correspondence problem makes it infeasible to increase the number of LEDs significantly and still expect real-time operation.

### 1.2. Object Tracking Using Deep Learning

About a decade ago, the price-performance ratio of computational hardware reached a critical level that enabled researchers to turn to complex machine learning (ML) methods for feature detection and tracking. Today, Convolutional Neural Networks of enormous complexity simulated on Graphics Processing Units (GPUs) are capable of identifying image features with complex appearances. In such a system the model is encoded in the weights between millions of simulated neurons. As building such complex models ‘by hand’ is virtually impossible, these weights are typically calculated by using backpropagation during a supervised learning phase in which training data samples demonstrate the desired mapping between inputs and outputs. Once trained, the network can perform inference, which is to take input data, process it through its neurons and provide outputs with—hopefully—expected results. These ML-based approaches are usually called Deep Learning (DL) methods due to the large number (depth) of computational layers. DL-based methods have been tremendously successful over the past few years to track both human [32, 33] and other animal [9] behavior. These methods dramatically cut down the number of training data frames required by transferring learning from a previously trained network to a new network [34].

While DL-based methods excel at recognizing features in images of realworld objects, current networks also face some challenges. Networks are only as good as their training data. In order to recognize a complex model, the training data needs to contain a balanced set of images representing diverse appearances. Animal behavioral training data may not contain some rarely observed configurations. The ground truth used to train neural networks is selected and often manually labeled by human experimenters—this leads to biases and inaccuracies in the training data. These 2D inaccuracies are often amplified when these positions are used in 3D pose estimation.

Vision-based 3D orientation estimation also poses challenges to DL-based approaches, due in large part to the topologically nontrivial nature of the space of 3D rotations, making, for example, Euler angles unsuitable near singularities [35]. Smooth, one-to-one representations as 3D submanifolds of 5D and 6D Euclidean space are more suitable for machine learning methods, but the accuracy of these systems is still significantly lower than marker-based methods [36, 37]. Hence, instead of directly estimating an object’s orientation using DL, most existing solutions use the neural network only for detecting features in 2D, and then employ classical methods to calculate the 3D pose from the positions of localized features. If the 3D geometry of the detected features is known, pose estimation is possible using a single camera (as in our method). However, if the feature geometry is not known, images from multiple cameras are required. Features are chosen to be camera-invariant, and thus the same network can be trained using frames from multiple cameras [38].

With the rising prominence of DeepLabCut for markerless tracking of laboratory animal features [9] such multi-camera 3D pose estimation systems based on DeepLabCut have been introduced [8, 38]. DeepLabCut-based systems can also be used to track features in real time [39, 40, 41], although processing speeds are severely limited by image resolution. While these algorithms can run using a CPU alone, their performance is degraded by up to 100 fold [38], rendering them too slow for real-time use.

In comparison to these methods, our requirements were to develop a precise monocular 3D pose tracker that does not require expensive GPU processing, can work at high frame rates and image resolutions, works for a wide range of viewing angles, and can incorporate a marker constellation with a large number of features.

### 1.3. Accurate, Robust, and Efficient Monocular 3D Tracking

Model-based systems can provide extremely accurate tracking by employing markers, while DL-based approaches are capable of directly tracking animal body parts with reasonable accuracy at the price of more costly hardware requirements, although real-time implementation has been achieved [39]. DL-based systems can provide sufficient accuracy and extraordinary tracking robustness for 2D tracking. However, for 3D tracking these systems might not be the best choice due to their potentially limited accuracy. This is particularly true for monocular systems where sub-pixel accurate feature detection accuracy is vital for accurate model-based orientation recovery, as discussed below. While the detection of simple markers can be done to a very high precision using simple image processing methods, the same cannot be said of animal body part detection accuracy using DL-methods.

Monocular 3D pose estimation is extremely sensitive to observation errors, especially when a limited number of features are available [42]. Such is the case when the target is small and can only accommodate a small number of identifiable features (this is certainly true when tracking small animals). In this work, we present a monocular 3D tracking system for small animal tracking that consists of a compact, lightweight visual target comprising a set of retro-reflective markers, a camera equipped with a ring-light, and a personal computer. The proposed system enables high-accuracy 3D tracking over a significantly wider range of view angles than other single camera systems in the literature. Marker localization is performed using model-based image processing methods and 3D pose estimation is done using a PnP algorithm [43, 44, 45].

### 1.4. Contributions

The primary contributions of the system presented in this paper are (1) the custom visual target design which enables the calculation of pose estimate from almost any orientation and (2) a computer vision algorithm that is capable of solving the difficult correspondence problem between markers and observations in real time. These two components—marker design and vision algorithm—were designed in concert.

The reliability of the proposed system was evaluated based on several hours of rat head tracking experiments. We validated its accuracy using a commercial wide-baseline multi-camera optical tracker, and compared its performance to the ArUco ARTag tracking solution. Finally, we demonstrated the use of the tracking system during neurophysiological recordings.

### 1.5. Terminology

We use the following terminologies to refer to objects that are handled by the tracking system:

#### Marker

Retroreflective sphere or hemisphere that appears as a bright spot on the image of the camera when illuminated by a strong ring light mounted on the camera’s lens.

#### Visual target/Target

Complex 3D object that is not rotationally symmetric. Our visual target comprises an assembly of markers mounted on a lightweight plastic frame that is sufficient to enable the unique determination of the pose of the target as it is tracked by a monocular tracking system.

#### Observation

Small bright spot detected on the camera’s image representing a marker candidate. This spot might represent an image artifact (for example lens flare) or the reflection of a small glossy object other than a marker.

## 2. Materials and Methods

The hardware components of the proposed tracking system consist of a marker, a camera, a ring light mounted on the camera, and a computer. During operation, the computer captures images from the camera, processes them, and generates the 3D position and 3D orientation of the marker for each video frame with respect to a specified reference frame. The system can also process video recordings offline.

### 2.1. Design Considerations

Our goal was to develop an affordable wide-angle 3D tracking solution for small animal research that can be deployed with minimal effort on hardware commonly available in behavioral laboratories. Experiments often involve the construction of a test environment in which animals are placed, and the environment may need to be equipped with sensors and actuators. Electrophysiological or imaging equipment may also need to be deployed. Multi-camera tracking systems are expensive, require complex calibration and setup process, and need to share space with other, often bulky, equipment.

In order to enable 3D pose reconstruction from a single camera view, we designed a visual target that enables visibility from a wide range of perspectives and a corresponding software system that is able to identify its unique pose. The visual target is lightweight and small in order to prevent it from interfering with behavior but also strong enough to maintain structural integrity despite any impacts that it might sustain during behavior. For our purposes, it accommodates electrophysiology equipment when mounted on the head of a rat. For affordability and ease-of-manufacture, the visual target can be manufactured with a 3D printer based on open source 3D designs; the designs are also easily customizable for different scales or to accommodate different equipment, viewing angles, etc.

The system uses IR illumination, leaving the experimenter the freedom to employ whatever visible lighting condition is required for the purpose of the experiment. We also required that the tracking algorithm be able to fall back to 2D tracking when there is not enough information on the image to resolve accurate 3D pose, as could happen for example in behaviors such as grooming that introduce partial occlusions of the visual target.

Our system operates in real time at a high frame rate on a personal computer equipped with a mainstream, 6-8 core CPU. It uses an open-source GNU/Linux operating system and our own custom, open-source tracking software. System calibration and setup software enables end-users to build and run the system without further assistance. Finally, the system can store its results in an accessible, open-source format, communicate with other equipment involved in the experiment, and synchronize the tracking results with the rest of the apparatus.

### 2.2. Visual Target

Our custom visual target enables tracking in a wide range of orientations, yet its small size and lightweight construction allow it to be mounted on small animals (Fig. 2). The target structure is modularized with an external framework for the optical markers, which then mates with adapters specific to different electrophysiology headstages. Adapters have been designed to house the NeuraLynx (Neuralynx, Bozeman, MT) Quickclip headstage, the FreeLynx wireless acquisition system, and the SpikeGadgets (Spikegadgets, San Francisco, CA) HH128 headstage. The external framework has an opening on the top to enable FreeLynx battery replacement without disassembly. The target consists of a 3D printed plastic ‘globe’ (with a diameter of 54 mm and a height of 39 mm) that has 16 sockets on its surface for holding retroreflective markers. Three of the markers are spheres of 7.9 mm diameter (size A) and the other 13 are hemispheres with a diameter of 3 mm (size B). Having retroreflective markers in two different sizes facilitate more efficient 3D pose estimation. The visual target weighs 3 g without and 3.5 g with the markers. The 3D printable CAD model of the target and instructions for assembly will be made publicly available online.

**Figure 2:**
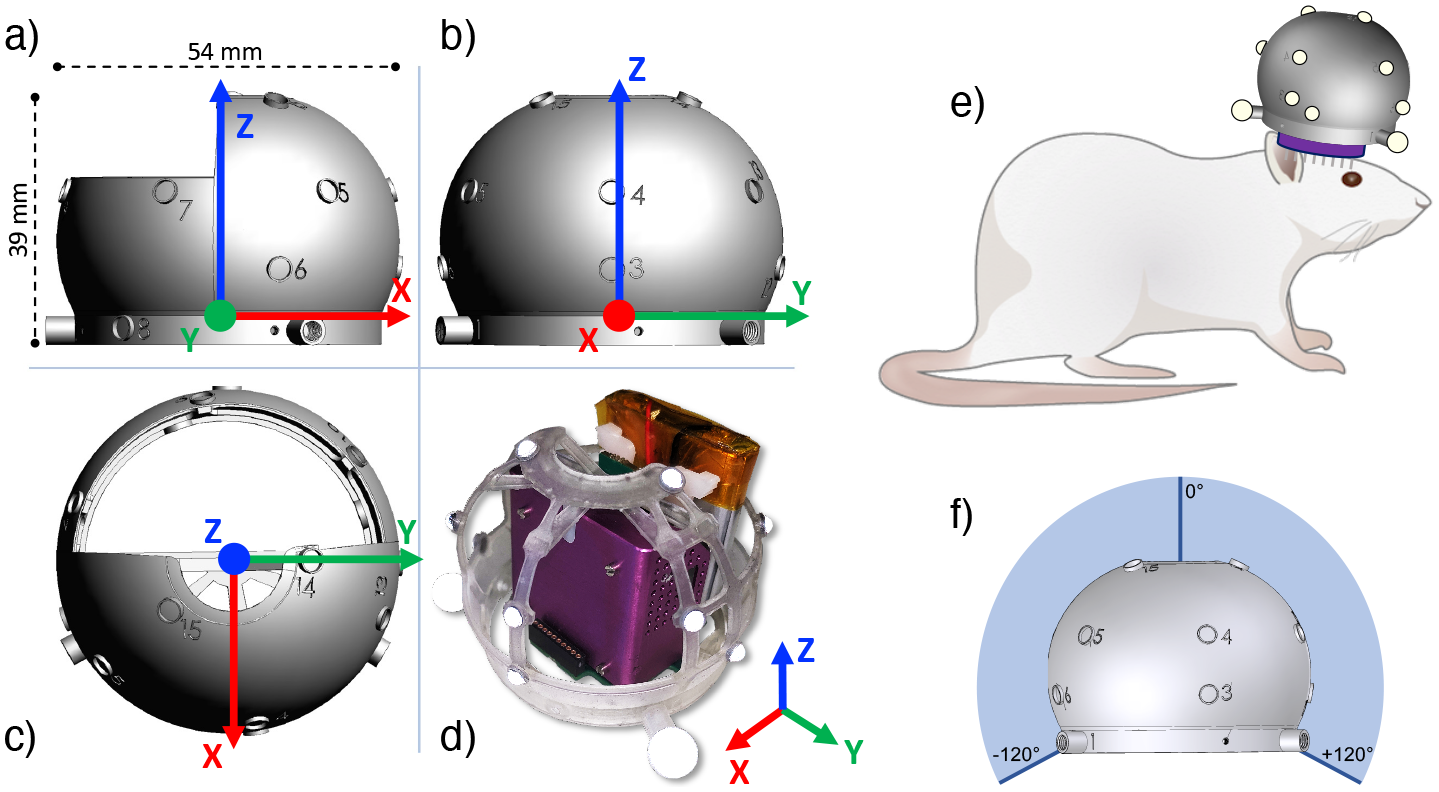
Visual target design. **(a-c)** Orthographic renderings of the target’s CAD model. The visual target measures 54 x 54 x 39 mm and weights 3.5 g. It features 16 spherical or hemispherical retroreflective markers. **(d)** Close-up photo showing the retroreflective markers mounted on the target’s clear plastic frame that houses the NeuraLynx FreeLynx wireless acquisition device (purple) and battery (copper). **(e)** Illustration of the target mounted on the head of a rat. **(f)** The markers are arranged in a configuration that enables full 3D (i.e. 6 degree-of-freedom position and orientation) tracking, enabling full 360^◦^ rotation around the nominal *z*-axis, and up to approximately ±120^◦^ range around the nominal *x* and *y* axes. This allows an animal to fully explore an environment with substantial fore–aft (“pitching”) and side-to-side (“rolling”) of the head, without losing 3D reconstruction.

The locations of the 16 retroreflective markers have been optimized so that at least six are always visible from any direction within the range of supported orientations, and that the geometrical configurations of visible markers are always unique. This enables a suitable algorithm to calculate the rotation of the target corresponding to any supported physical orientation.

In reference to the coordinate frame depicted in Fig. 2, the visual target can be observed so long as the *z*-axis is rotated no more than 120^◦^ relative to the line of sight from the camera, i.e. the *z*-axis can cover more than an entire hemisphere. This range is typically sufficient to track the head of a small animal during a foraging and other behaviors where the animal is generally oriented upright.

### 2.3. Tracking Method

The tracking algorithm was designed to track the visual target by first locating the bright spots representing reflective markers on camera images (observations) and then resolving the observation-marker correspondence and the 3D pose of the target in a combined optimization framework. The target’s 16 markers were arranged in a geometry such that the projection of those markers is unique from any point of view; therefore, the tracking algorithm will find a single unique solution for any given pose of the visual target.

The method consists of three main steps: marker detection on video frames, spatiotemporal tracking of the target, and localization of the visual target by solving the correspondence problem (Figs. 3, 4). An optional initialization step – anchor set optimization – increases reliability by automatically finding the strongest visual features available on the target and optimizing the localization process for the detection of those features. The prerequisites of accurate and robust tracking are camera intrinsic calibration and proper camera exposure and focus, which are described in detail in the System Calibration and Setup2.5 subsection.

**Figure 3:**
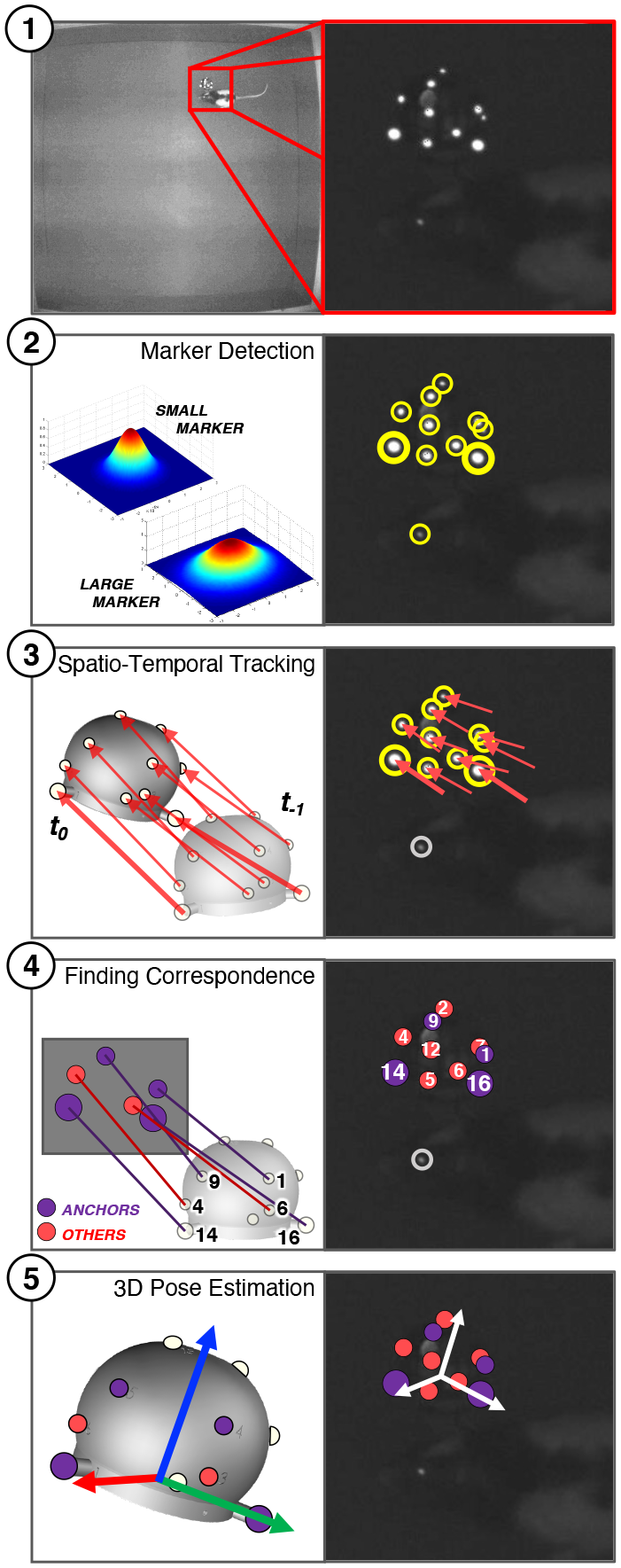
Illustration of the processing steps of the 3D tracking method: **(1)** Finding the region-of-interest (ROI) on the grayscale image using connected component analysis and clustering, then dewarping the ROI to eliminate local camera distortions; **(2)** Sub-pixel-accurate estimation of marker positions and recognition of marker sizes; **(3)** Attempting to find marker correspondence by matching new marker detections to marker positions predicted from past trajectory (spatiotemporal tracking); **(4)** Performing combinatorial correspondence matching when spatiotemporal tracking (step 3) fails to determine correspondence; **(5)** Calculating final 3D pose from correspondences.

**Figure 4:**
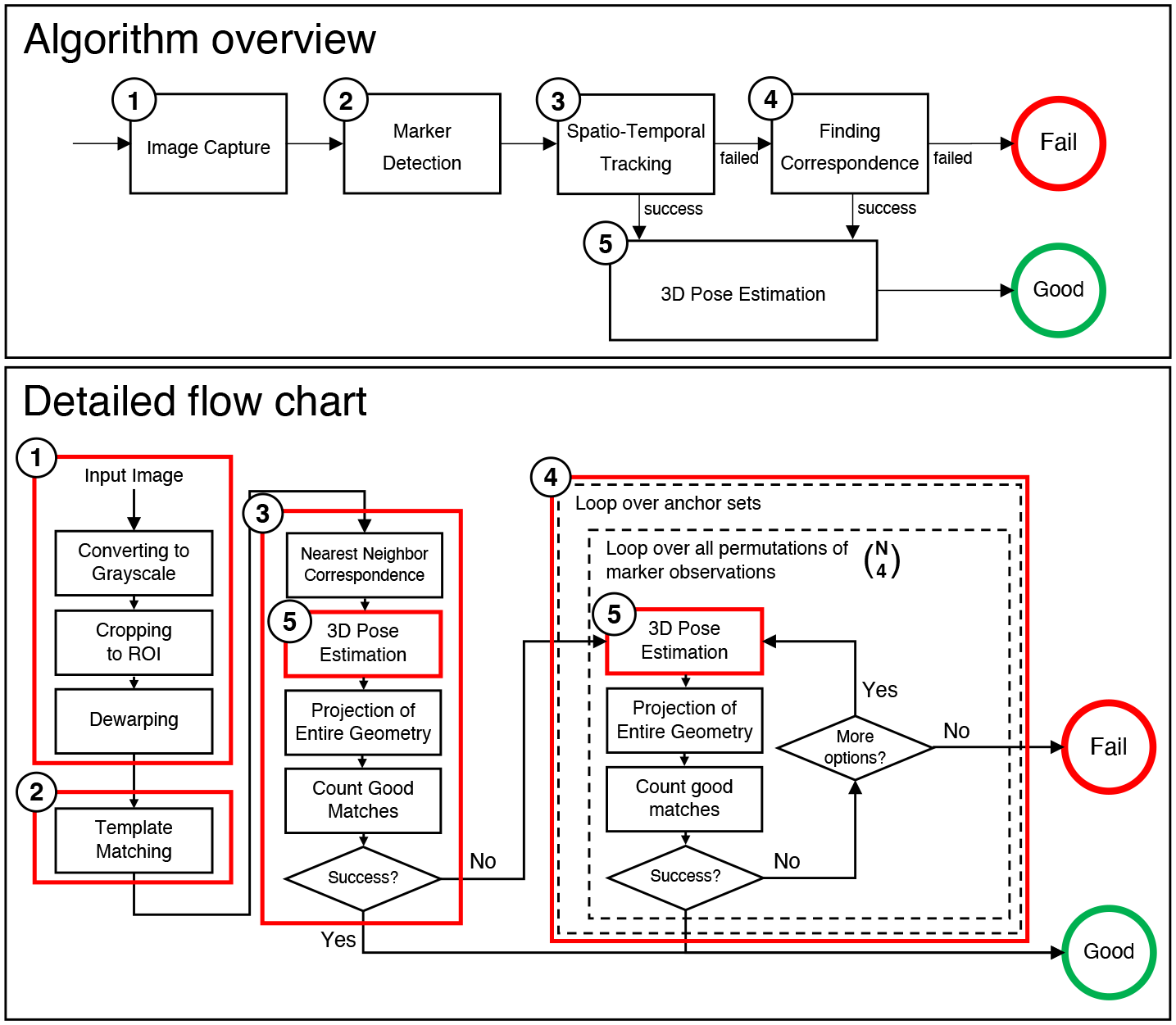
Processing steps of the proposed 3D tracking method: **(1)** Image capture and dewarping of region-of-interest (ROI); **(2)** Marker detection; **(3)** Attempting to find marker correspondences using spatiotemporal tracking; **(4)** Combinatorial correspondence matching when spatiotemporal tracking has failed; **(5)** Calculating the 3D pose from marker positions and correspondences. **Top:** Overview of processing steps; **Bottom:** Flow chart illustrating sub-processing steps and the combinatorial phase.

#### 2.3.1. Marker Detection

Tracking the visual target on any given frame starts with simple image processing steps to locate a cluster of small bright spots on the image.

First, the region-of-interest (ROI), defined by a bounding rectangle on the input video frame, is dewarped to eliminate radial distortion. The dewarping process requires camera intrinsic parameters determined during offline camera calibration. The ROI covers the area where the visual target is expected to appear on the image. Initially, the ROI covers the the entire image, but after the first successful detection of the target, the ROI is narrowed down to a small neighborhood of the target’s image position. The position of the ROI on new video frames is predicted based on the recent velocity of the target. If the tracker fails to locate the target, the ROI is gradually expanded until the target is located or eventually the ROI encompasses the entire image. When the target is successfully localized again, the ROI shrinks again to the narrow neighborhood around the target.

The target appears in the ROI as a cluster of small bright spots. While most spots represent individual markers, some might be bright or shiny objects that are not part of the target. The marker detection algorithm identifies small bright spots by matching template images to the ROI. The number of templates is determined by how many unique sizes of markers are used in the visual target. The templates are 2D Gaussian functions generated to match the expected size of markers. The results of template matching are evaluated using normalized cross correlation (NCC), which is relatively insensitive to brightness and contrast variations. Bright spots that are dissimilar to the 2D Gaussian profile are discarded by template matching. Additional filtering steps eliminate markers that are outside a specified intensity range.

Once the initial collection of candidate markers is identified, the detection algorithm uses a clustering method to locate a single tight cluster among them. Candidates outside of the cluster are discarded as they are unlikely to be part of the target. Positions of spots are so far defined as pixel locations, which are only rough estimates of their actual positions. For sub-pixel accurate positions, the algorithm resamples the image of each candidate at 4x resolution using a 2D 4-lobed Lanczos kernel [46], and re-runs NCC-based template matching with 4x larger 2D Gaussian templates. The resulting matches on the oversampled image represent 0.25 pixel accuracy on the original image. Due to image noise and limited image resolution, resampling at even higher resolution does not seem to result in higher position accuracy.

#### 2.3.2. Monocular Pose Estimation and the Correspondence Problem

The 3D pose of a 3D point cloud can be unambiguously calculated from its 2D projection if there are at least four non-co-linear points in the point cloud and the correspondence between the projections and the 3D points is known [47]. The quality of the resulting pose estimate can be measured by using the the estimated pose to reproject the point cloud onto the image plane and measuring the distance (reprojection error) between the reprojected point coordinates and the original projections.

If the correspondence between projections and the points is not known, an algorithm may generate a list of potential correspondences and test them by measuring the reprojection error. However, the process of perspective projection from 3D to 2D reduces the dimensionality of the data and enables configurations where multiple different 3D point clouds projected from different 3D poses yield identical or very similar 2D projections. The chances of this ambiguity is particularly high when there are only four points in the point cloud, in which case it is likely that there are multiple potential correspondences with low reprojection error, making it impossible to find the correct correspondence. Increasing the number of points to five in the point cloud can eliminate or drastically reduce the chances of multiple correspondences with low reprojection error if the 3D configuration of the points is chosen suitably, for example by avoiding symmetries.

In our system we set the minimum number of matched markers to six in order to minimize the chance of ambiguous correspondences, and markers on the visual target were mounted in a geometrical configuration that reduces potential ambiguities.

#### 2.3.3. Solving the Correspondence Problem

The marker detection method provides a list of marker observations without further hints on their correspondence to physical markers. When the appearance of markers are indistinguishable from each other, correspondence between the detected spots on images and the points in the point cloud can be established in multiple ways. The number of possible correspondences is characterized by the permutation 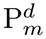, with

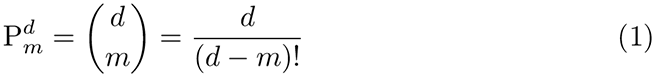

where *m* is the number of markers in the optical target and *d* is the number of detected spots on the image, i.e. the observations. 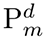 is relatively low if there are few markers and few observations but increases faster than exponentially when the number of markers and observations grow. For instance, for 10 observations and 6 markers (a common use case), the number of possible correspondences is 151, 200—a brute force method would struggle to process these correspondences in real time at a high frame rate.

Knowing the 3D geometry of the markers and the optical properties of the camera can be used to decrease the number of possible correspondences. We implemented three improvements for reducing combinatorial complexity: (1) we evaluate all possible correspondences only for 4 special markers (termed anchors) and rely on the known geometry of the remaining markers to select the correct correspondence, (2) we use markers of multiple sizes, which enables the marker detector to separate observations into multiple groups, and (3) we introduce simple geometrical constraints on how the markers are expected to be projected onto images, enabling fast filtering of invalid configurations.

##### Anchors

Introducing anchors reduces the number of comparisons in our example by a factor of 30, from 151, 200 to 5, 040 (*m* = 4 in Equation 1). There are ≥ 6 markers visible from any orientation, and at any particular orientation 4 of the visible markers are designated as an anchor set. To find the correct correspondence between markers and observations, the algorithm first calculates— for each possible anchor correspondence—the 3D pose of the target (using the Perspective-n-Point (PnP) algorithm), which then enables the prediction of the positions for the remaining visible non-anchor markers. The predicted marker positions are matched with the observations on the image using the nearest neighbor method under the 2D Euclidean distance metric. The configuration with the highest number of non-anchor markers that match the observations is then selected as the correct correspondence. Once the correspondence for all the matching observations is established, the 3D pose is refined by another run of the PnP algorithm using all the matched observations that results in a more accurate measurement than the initial estimate based on only 4 anchors.

While using anchors reduces the combinatorial complexity significantly, the selection of these 4 anchors limits the range of directions from which the target can be tracked. Our target has 16 markers with 6 guaranteed to be visible from any particular direction—therefore an anchor set will only be detectable when the target is oriented such that all the anchors in the set are visible. To solve this, the tracking method uses multiple anchor sets, each representing a limited range of orientations from which the target is observed. The combination of all anchor sets completely cover the supported range of orientations. With this, the maximum number of comparisons is reduced to 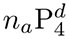, where *n_*a*_* is the number of anchor sets.

We sort the list of anchor sets between frames to reduce the actual number of correspondence comparisons even further. The tracker performs correspondence comparisons sequentially from the list of *n_*a*_* anchor sets. Once an anchor set is found, the tracker tries that anchor set first for consecutive video frames. When the orientation of the target changes so much that the initial anchor set is not fully visible anymore, the method will sequentially proceed through the list. The list of anchor sets is periodically re-sorted in descending order of utilization. The search thus starts with the most likely-to-succeed anchor sets, based on recent usage statistics.

##### Different size markers

The method supports visual targets featuring multiple marker sizes. During the marker detection phase, bright spots are classified to one of the size classes based on their matches to the different size 2D Gaussian kernels. In the correspondence phase, each marker is only matched to observations in its own size class, thereby reducing the number of correspondence comparisons by a significant amount. For instance, with 2 types of markers (large and small), 2 large and 8 small marker observations, and one anchor set featuring 1 large and 3 small markers, the number of comparisons is 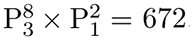. If marker sizes are ignored, the number of comparisons is 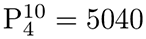

##### Filtering configurations

This improvement takes advantage of the properties of perspective projection to eliminate certain anchor observation configurations from correspondence comparison. When four 3D points that define the corners of a polygon are projected using perspective projection to a 2D surface, certain properties are preserved such as convexity and whether the points of a convex quadrilateral are defined in clockwise or counterclockwise direction. Testing these properties of a four sided polygon in 2D is simple and efficient. The tracker algorithm requires that anchor sets be defined as sets of 4 anchors that are approximately on the same plane, that they define a convex shape on their plane and that the corners are defined in clockwise order when observed from the visible side. When these requirements are met, the camera projection of the visible anchor sets also describe a convex 2D polygon with its corners defined in clockwise order. In the correspondence comparison the algorithm checks if any given set of 4 observations that are to be matched with 4 anchors meet these requirements before attempting to perform pose estimation. Configurations not meeting the criteria are discarded, which cuts the number of fully evaluated configurations by a factor of 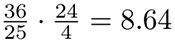, as the probability of four randomly picked points on a rectangular plane to form a convex polygon is 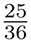 [48], and 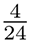 ordered sequences of four such points are clockwise.

#### 2.3.4. Spatiotemporal Tracking

The correspondence phase is capable of determining the pose of the optical target on individual frames without having any knowledge of the pose of the target on preceding video frames. In behavioral tracking, the pose of the target can be expected to correlate with its pose on preceding frames. If the frame rate is adequately high compared to the rate of motion of the target, the pose changes between consecutive video frames can be estimated using spatiotemporal tracking techniques. The tracking method assumes that the position and orientation of the target changes smoothly in time and therefore their values can be predicted with reasonable accuracy at least one frame time ahead. When the tracker was able to successfully determine the pose of the target for at least two consecutive frames, it calculates the angular and translational velocities of the target based on these frames and predicts the pose for the next video frame by assuming constant velocity. When the next video frame arrives, it detects the positions of bright spots on the image in the neighborhood of the predicted position, and matches the observations to the predicted marker positions using the nearest neighbor method. Once a correspondence is established, it is tested for validity using the PnP algorithm. If the resulting 3D pose is near the predicted pose, the new pose is accepted and the tracker skips the time-consuming correspondence computations.

#### 2.3.5. Anchor Set Optimization

The selection of suitable anchor sets is essential for efficient tracking. Anchor sets form the foundation of an efficient combinatorial solution to the point correspondence problem. The tracking algorithm requires enough anchor sets to cover all supported orientations, but using too many anchor sets slows down processing. Finding the right balance between the number of anchor sets and the anchor configurations inside those anchor sets is required for efficient tracking. There are billions of possible anchor set configurations that cover the desired orientations but only a few of these configurations combine robust tracking with a low number of anchor sets. Initially, we selected the list of anchor sets manually by visually inspecting the target from every orientation and taking notes of suitable looking anchor sets. The process worked reasonably well but in our evaluations we failed to achieve better than 90% detection success rate.

To overcome this, we developed an algorithm to optimize the set of anchor sets for a given optical target. The algorithm has three main steps: simulation, optimization, and minimization.

##### Simulation

The algorithm first generates a large number (by default 10,000) of random projections of the optical target with uniform distribution in a specified range of orientations. To sample orientations uniformly, we ignored the symmetry around the optical axis of the camera, which allowed us to sample uniformly over a simple sphere. In each simulated view, the algorithm iterates through all possible permutations of four detected markers and selects the ones that satisfy the requirements (convexity, vertices defined in clockwise order) for anchor sets and contain at least two different types (sizes) of markers. The anchor sets are hashed in a 32-bit unsigned integer. Anchor sets are considered identical by the hash function if they are cyclic permutations (for example 312-14-7 and 7-3-12-14 are identical). For each view, the algorithm saves the list of anchor-set-hash values, that represent the anchor set candidates visible from the view.

##### Optimization

The algorithm selects a few robustly detectable anchor sets from the candidates identified by simulation that, taken together, cover the entire range of supported orientations.

The optimization algorithm is formulated as a minimum spanning tree problem. The anchor set candidates, their simulated views, and their relationships are represented in a weighted graph as shown in Fig. 5a. Anchor sets are represented by nodes and labeled by their hash values (nodes labeled A-D in the figure). Simulated views are also represented by nodes and labeled by numbers (1–number of views). Edges between anchor sets and nodes represent the visibility of anchor sets from simulated views. An edge between an anchor set and a view is weighted by the reciprocal of the number of views visible from the anchor set. If the anchor set has *N* views associated with it, then the edges connecting views to it will all be weighted 1*/N*. There is one additional root node (labeled *X* in the figure), that is connected to every anchor set by an edge with a weight of zero.

**Figure 5:**
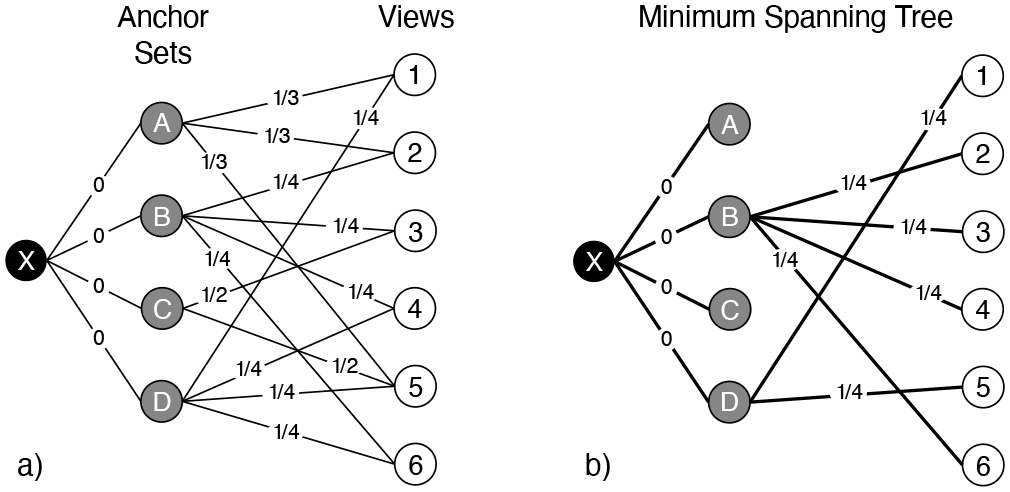
Illustration of the anchor set optimization algorithm. **(a)** Anchor set candidates (labeled A-D) and views (labeled 1-6) are calculated by simulating allowed view angles. The real-world graph for our visual target is significantly larger than this illustration, containing 10,000 simulated views and ∼3,000 anchor set candidates. **(b)** The first step of the anchor set optimization method uses the minimum spanning tree (MST) algorithm to find the best anchor sets that cover the entire range of supported orientations. This solution is not necessarily unique and may have redundancies, therefore a second processing step (not shown here) is required to minimize the number of anchor sets.

The minimum spanning tree (MST) of this graph has some useful properties (Fig. 5b). The MST will have exactly one edge connecting each view to one of the anchor sets. For each view the MST will keep only the edge that connects it to the anchor set that has the most associated views. By minimizing the sum of weights of the edges in the spanning tree, the MST will favor a configuration where the views are connected to highly visible anchor sets, while anchor sets with lower visibility tend not to be connected to any views.

The result of this optimization is the list of anchor sets that are connected to at least one view. For our target, the optimization selects 16-18 anchor sets out of the several thousands. The variability in the number of anchor sets is due to the stochastic nature of the simulation. In our experiments the optimized anchor sets provide highly robust operation that reduces failure rates by a factor of ∼20x compared to manually selected anchor sets.

##### Minimization

This step is optional, as it reduces complexity and achieves higher frame rate at the potential expense of slightly degraded tracking reliability. Running the tracker with minimization enabled is designated as FAST mode. While the anchor sets selected by the MST are highly robust due to their high visibility, the number of selected anchor sets is not minimal. The MST has multiple solutions that result in the same minimum weight and the solver selects one unspecified instance of them. However some of the solutions would be preferred over others in our application, as we also aim to reduce the number of selected anchors to improve combinatorial performance.

The ultimate minimal solution requires an additional step of post processing, and it reduces the number of anchors to 9 (for our specific target), approximately half the size of the simple MST solution. This minimal solution is also guaranteed to cover all simulated views and prioritize high visibility anchor sets; however, it will have fewer redundancies (overlaps between regions covered by anchor sets), and therefore it will be somewhat less robust than the simple MST solution.

The minimization algorithm first sorts the anchor sets in descending order based on how many views they are associated with. Starting from the anchor sets with the lowest visibility, it then examines each one if they are redundant. An anchor set is redundant if all of the views to which it is associated can be accessed through other selected anchor sets. Each anchor set that is found redundant is removed from the list of selected anchor sets.

### 2.4. Software and System Architecture

#### 2.4.1. Online Processing

The tracking system runs on Ubuntu Linux and uses the Robot Operating System (ROS) software framework. ROS is an open source software that features robust support for a wide range of cameras, data recording and playback capabilities, and provides means for communication with other software components of the experimental apparatus.

Using ROS enabled us to make live tracking results accessible by external software in a flexible way. Researchers intending to integrate the tracker functionality into their closed-loop experiments only need to implement a small ROS software interface between their own code and the tracker. For this software interface, ROS supports C++, Python, and Matlab implementations. ROS also has the convenient capability to make the interface work locally (i.e. between software programs running on the same computer) and remotely (i.e. tracking software running on a separate computer, accessible through local network). While our tracker code was designed to run best on Linux, the external software interfacing with it can run on Linux, Windows, or MacOS [49].

#### 2.4.2. Data Recording and Offline Processing

ROS has built-in data recording and playback functionality. ROS *bag files* are universal containers for storing one or more simultaneous timestamped ROS data streams, including video data and tracking results. Video data recorded into *bag files* can be processed offline by the tracker and the software makes sure that video timestamps are carried over to the corresponding tracking results.

#### 2.4.3. Time Synchronization

Data records transmitted through ROS *topics* are always timestamped by the publishing *node*. Timestamps are defined in Coordinated Universal Time (UTC). In the case of *image topics* containing live video, the timestamps assigned to video frames are generated by the camera capture *node* at acquisition time. For data in other topics, it is the responsibility of the original publisher to generate accurate timestamps. Topics that include data that was not published by the original source but contain secondary, processed data typically inherit the timestamp of the original data source. In the case of the tracking software, the timestamp of a tracking result will inherit the value of the original timestamp of the video frame to which it belongs.

If the tracking software is used in an apparatus that has multiple types of computing hardware, each with a separate clock, the clocks between the computers need to be synchronized before running the tracking software. For synchronizing the clocks between multiple computers, we recommend configuring the operating system of each computer for automatic synchronization to the same time server using the Network Time Protocol (NTP) or Precision Time Protocol (PTP).

Another option is available when working with a neural recording system, which generally is able to accurately timestamp TTL pulses. The tracking computer is equipped with a DAQ and set to generate a randomized TTL pulse train (mean 10 s between pulses, 1 s pulse duration) that is fed into the digital inputs of the neural data acquisition system. The paired timestamps of these pulses are then used post-hoc to synchronize the neural and experimental data streams using the Needleman-Wunsch algorithm [50, 51].

#### 2.4.4. Hardware Requirements

The current version of the software was designed for the Ubuntu Linux 18.04 (or newer) 64-bit operating system and a compatible computer. The tracking software is capable of using up to 8 threads for processing; therefore a CPU with at least 8 CPU cores is required for best performance. The frame rate of tracking is variable, as different tracking modes have different computational complexities. The lowest frame rates are expected at the worst case scenarios when the tracker fails to resolve marker correspondence. To improve frame rate in these situations, a CPU with an operating frequency of 3 GHz or better is recommended.

The resolution and frame rate of the camera significantly affect the reliability of tracking. Image resolution needs to be high enough that small markers in the visual target appear at least 4 pixels in diameter. For tracking the rapid motions of a small rodent, a camera with a frame rate of at least 45 fps (frames per second) is recommended.

#### 2.4.5. Access to the Software

The tracking software will be made publicly available on GitHub for free under the MIT License [52]. The software package includes a user’s manual and 3D printable CAD drawings of the visual target with assembly instructions.

### 2.5. System Calibration and Setup

#### 2.5.1. Camera Optical Calibration

Before accurate geometric measurements can be made based on observations of the visual target on the camera image, the exact optical properties of the camera and its lens need to be measured and stored in a configuration file. These properties are described by two sets of intrinsic parameters. The raw intrinsic parameters contain the focal length, the position of the optical center, and the five parameters of radial distortion, all of which need to be measured by an offline camera calibration method. The undistorted intrinsic parameters contain the desired focal length and optical center position, which define the geometry of the undistorted image that the tracking software can use for processing. The two sets of parameters together enable the mapping of each raw image pixel onto an image with perfect perspective projection that is required for easy geometrical calculations.

There are several free and commercial camera calibration tools available, most of which can be used for determining the required raw intrinsic parameters, such as the Camera Calibration Toolbox for Matlab [53].

#### 2.5.2. Setup of Illumination and Camera Exposure

The tracker software is designed to localize small bright spots on the camera image. The retroreflective markers on the target appear as bright spots as long as the illumination is appropriate and the exposure parameters are correctly set for the camera. In order to minimize the brightness of other objects and the environment in the field of view, the intensity of illumination from the ring light mounted around the lens needs to be high enough so that the brightness of the markers appear significantly brighter than other features. Other reflective objects and light sources in the view may interfere with tracking performance and must be removed from the environment.

Camera exposure needs to be set to manual mode and the shutter speed needs to be increased until the bright spots representing the markers stop being saturated. Saturation is characterized by a flat white appearance; therefore, the shutter speed needs to be increased until the spots appear to have a spherical brightness profile with darker shades around the edges and a bright peak in the middle. Shorter exposure times (faster shutter speeds) also reduce motion blur in the images, which may significantly improve tracking reliability. We have found that, for our application, good tracking performance requires that the exposure time be under 2 ms.

#### 2.5.3. Setup of Visual Target and Scene Geometry

The visual target is defined as a point cloud in 3D space with each marker represented by a point. The description of each point includes its 3D coordinate, size class and visibility angle. The coordinates can be obtained from the CAD model of the target or estimated from multiple 2D images through image processing methods such as bundle adjustment. In our visual target, two different sizes of markers are used. The visibility angle specifies the range of angles from which the marker is visible. The small marker types on the visual target are hemispherical therefore their view angle is more limited than the large markers which are spheres.

Users may want to capture 3D tracking results with respect to a reference frame that is different from the camera’s coordinate frame. For example a particular point and orientation in the animal’s test environment may be used to define the reference frame. To facilitate this, the tracking software allows the user to specify the position and orientation of an optional reference frame in the configuration file.

#### 2.5.4. Animal Experiments

We performed behavioral and neurophysiological recordings using LongEvans rats (Envigo Harlan). All animal care and housing procedures complied with National Institutes of Health guidelines and followed protocols approved by the Institutional Animal Care and Use Committee at Johns Hopkins University.

## 3. Results

We performed both accuracy and reliability evaluation of the proposed tracking solution. For verifying accuracy, we compared our 3D tracker’s results to that of a surgical-grade commercial optical tracking solution. Reliability was evaluated by mounting the visual target on a rat’s head, tracking the animal while it was roaming in an open arena, and analyzing the recorded tracking results. We also compared tracking performance to another open-source monocular 3D pose tracking method that uses ARTags. The Near IR camera used for evaluating our tracking system’s accuracy and reliability is a Grasshopper3 USB3 (GS3-U3-41C6NIR-C, Flir Systems Inc., OR, USA) with a resolution of 2048 x 2048 driven at the framerate of 45 frames per second.

### 3.1. Evaluation of Accuracy

Measuring the real-world accuracy of an optical tracking system is difficult and sometimes impossible, which is why it is rarely done in research publications. The difficulty lies in the generation of accurate ground truth. For getting ground truth, the motion of the object or animal would either need to be simultaneously tracked by other, more accurate means, or the motion would need to be generated by the experimenter, for example by moving the object with a robot along a known trajectory. We found it most practical in our setting to benchmark our system against a commercial tracking system with well known tracking performance.

The accuracy of the tracking system was measured by comparing its tracking results to pose data acquired by a Polaris P4 commercial optical tracker (Northern Digital Inc. (NDI), Ontario, Canada). The P4’s average accuracy is better than 0.25 mm (*<*0.5 mm in 95% confidence interval) [54], making it suitable for collecting ground truth data. The experimental setup is illustrated in Fig. 7. For the evaluation, the camera for our tracking system was mounted facing down, ∼200 cm from the visual target which was moved around by an operator along a random trajectory. Attached to the same visual target were three large retroreflective markers (NDI Spheres), that were simultaneously tracked by the Polaris optical tracker, placed at ∼120 cm from the visual target, looking at the scene at a ∼45^◦^angle compared to the angle of our camera. After data capture, we registered the two datasets to each other (aligned the 3D positions and orientations), then compared the positions and orientations calculated by our tracker to the ground truth captured by the Polaris. The measured tracking errors are visualized in Fig. 8, and Table 1 (non-scaled) breaks down errors by coordinate axes. We found that the average position error is highest in the Z (vertical) direction (9.75 mm) and lowest in the XY plane (4.84 mm).

**Figure 6:**
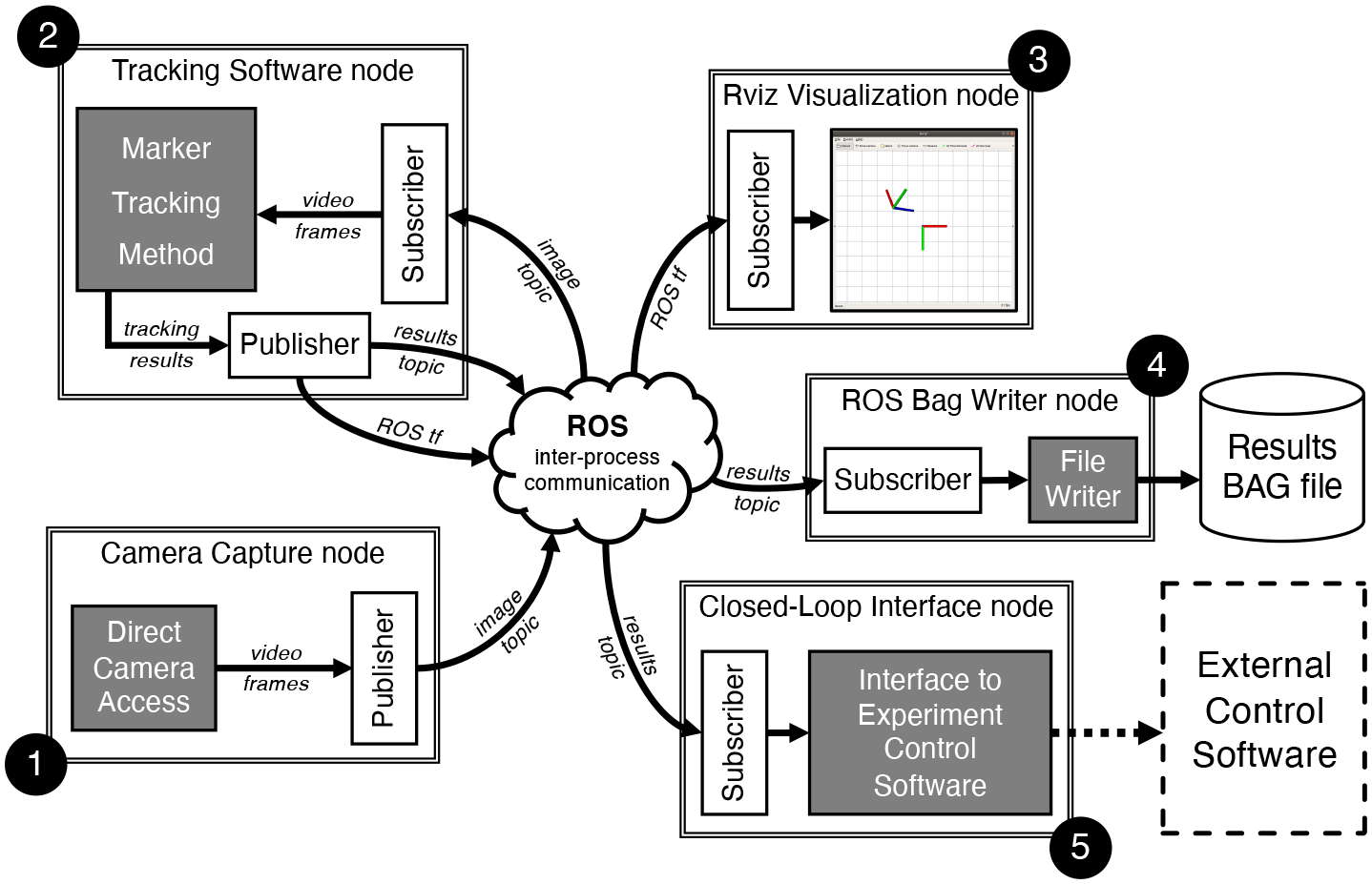
Tracking software architecture: The ROS software framework provides standardized inter-process communication channels (i.e. topics) between software components (ROS nodes). **(1)** Video capture from the camera is handled by the Camera Capture node, that publishes video frames in an image topic. **(2)** The Tracking Software node receives the video by subscribing to the image topic, processes video frames (i.e. detects the visual target on images), then publishes the tracking results on a topic and the 3D pose of the target in a special kind of topic (ROS tf) for visualization purposes. **(3)** Rviz is a visualization tool built into ROS that is capable of visualizing 3D coordinate frames published in the ROS tf topic. **(4)** Timestamped tracking results are recorded into ROS bag files by the ROS Bag Writer node. **(5)** An optional Closed-Loop Interface node may subscribe to tracking results and transmit them to an External Control Software. This node may be located on a separate computer connected to the tracking computer.

**Figure 7:**
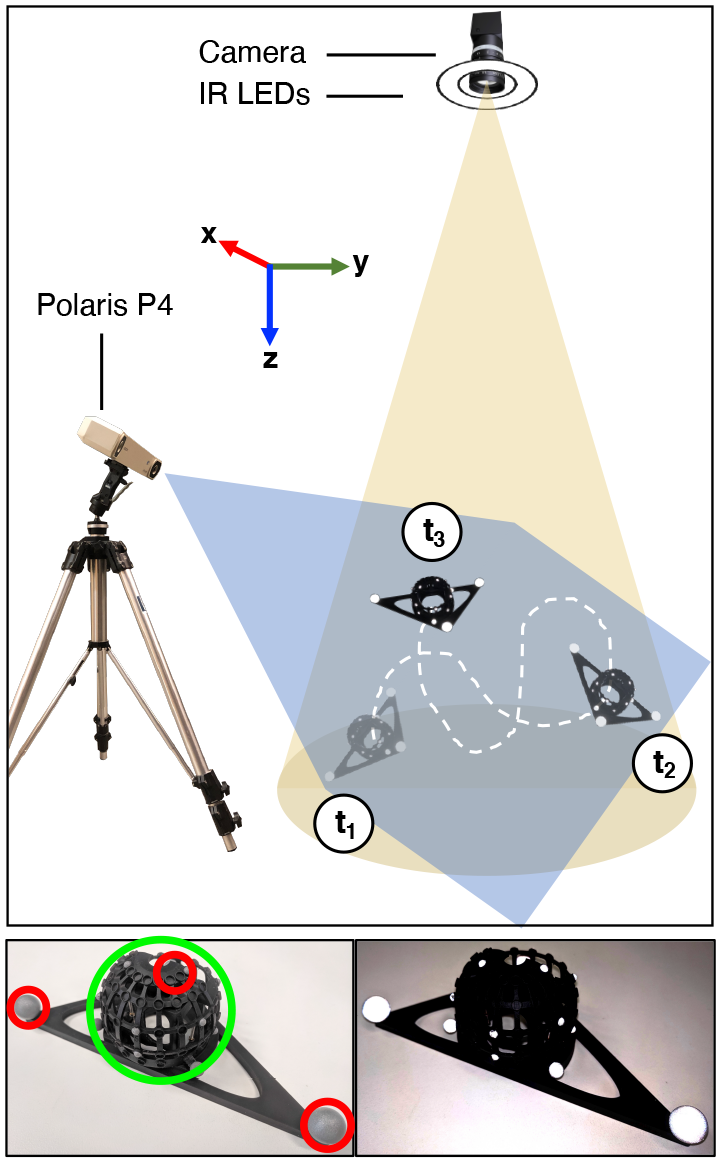
The accuracy of the proposed 3D tracker was measured using a surgical-grade, high-precision, wide-baseline (stereo) optical tracker (NDI’s Polaris P4). **Bottom:** The visual target (marked green) was rigidly mounted on a triangular frame that featured three large NDI retroreflective markers (marked red) in its corners. **Top:** During the evaluation, the P4 was tracking the locations of the three large marker from the side, while our monocular tracking system was tracking our visual target using the camera mounted above, at a distance of ∼2 m from the target. The target was moved by hand in a random trajectory (dashed white line) while making sure that all three large markers were always visible to the P4. Tracking data was recorded into a ROS bag file from both the Polaris and our tracker.

**Figure 8:**
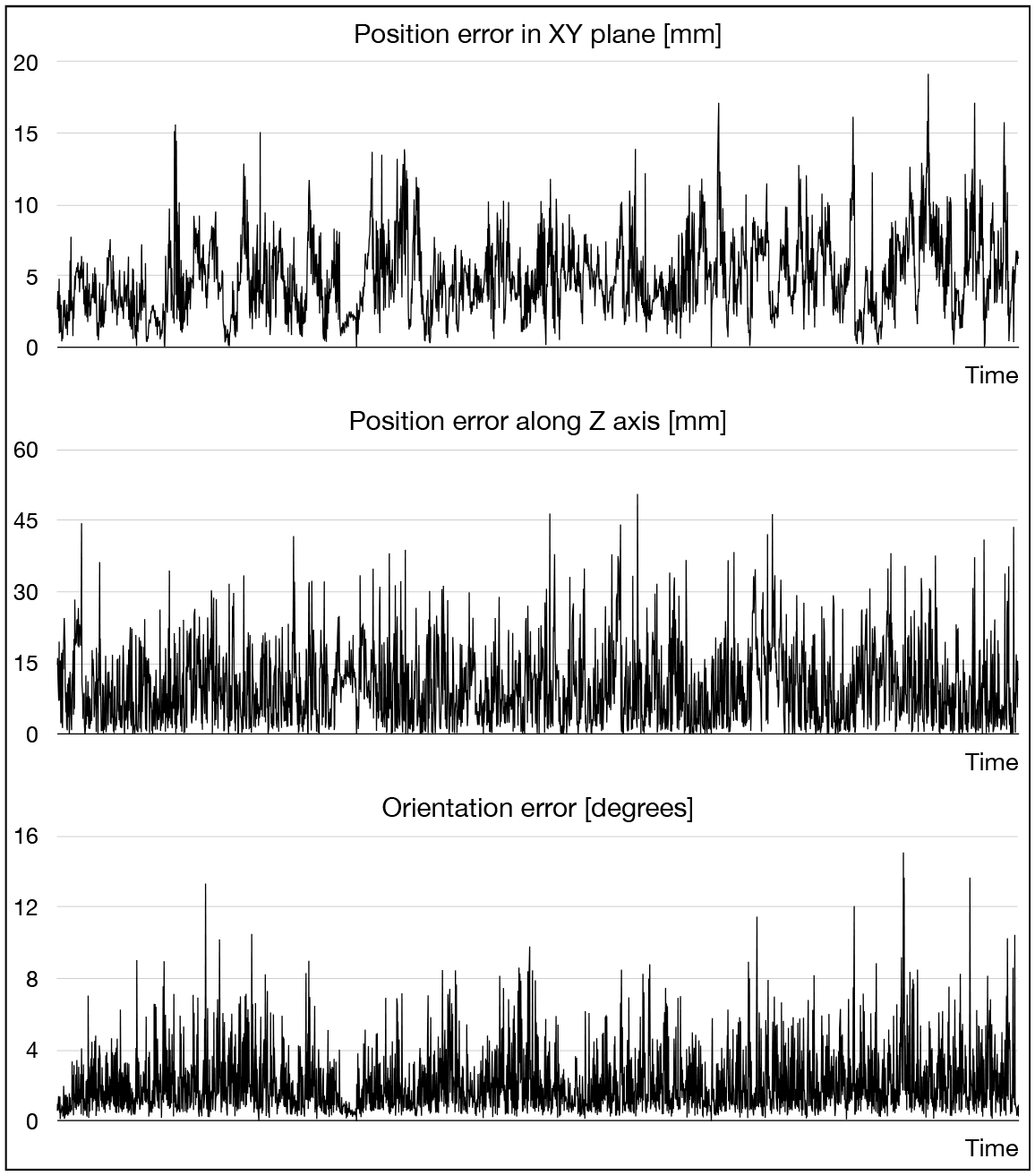
The position and orientation accuracy of the 3D tracker was measured with the help of a high precision wide-baseline commercial tracker (Polaris P4). The accuracy of groundtruth recorded by the P4 is known to be better than 0.25 mm. As the spatial relationship between the P4 and our camera was not known, the two trajectories recorded by the P4 and our tracker had to be registered to each other, which then enabled the calculation of errors. **Top-middle:** The position errors along the plane parallel to the image plane (XY) were lower than the errors measured along the camera axis (Z), which is expected due to the difficulty of accurately resolving distance from a single camera view. **Bottom:** Mean orientation error was around 2^◦^.

**Table 1:** Tracking accuracy compared to Polaris P4 optical tracker.

|  | Unit | Non-Scaled |  | Scaled |  |
| --- | --- | --- | --- | --- | --- |
|  |  | Mean | StDev | Mean | StDev |
| Position error (XYZ) | [mm] | 10.90 | 7.96 | 10.16 | 7.73 |
| Position error (XY) | [mm] | 4.84 | 2.51 | 3.30 | 2.20 |
| Position error (Z) | [mm] | 9.75 | 7.55 | 9.60 | 7.41 |
| Orientation error | [deg] | 1.96 | 1.56 | 1.96 | 1.56 |

In the camera’s coordinate frame, the Z coordinate can be interpreted as distance from the camera. The bulk of the position error is concentrated along the Z axis, which is due to relying on a single camera to resolve the visual target’s distance. The mean orientation error in full 3D rotation space was measured at 1.96^◦^.

During our analysis we noticed that the target trajectory calculated by the our tracker was scaled ∼2% larger than the trajectory provided by the Polaris. This consistent discrepancy is likely due to inaccuracy in camera intrinsic calibration. Using a camera focal length for 3D pose estimation estimate that is 2% off the actual value would result in the same trajectory scale difference. After correcting for the 2% scale factor difference, the accuracy of the tracker improved from 10.9 mm to 10.16 mm (Table 1-scaled).

The focal length—and other parameters—calculated during camera calibration are estimates, typically characterized by mean and variance. For 3D pose estimation the mean values are typically used; however, the mean only represents the best estimate within a range. Camera calibration uncertainties, manifest as high variance estimates, can be mitigated in a number of ways, such as increasing the number of calibration images or using larger calibration objects. Fortunately, an overall 2% uniform scaling of animal head trajectories would not likely change our overall interpretations of the types of behavioral data we are investigating, but this sensitivity to camera intrinsic calibration accuracy does highlight a disadvantage of monocular 3D tracking compared to multi-camera tracking methods.

### 3.2. Evaluation of Reliability Using Freely Behaving Animal

For determining the reliability of the proposed 3D tracker, we performed a series of laboratory experiments tracking the head of a rat moving freely in an open arena within the field of view of the camera at a distance of ∼210 cm, as shown in Fig. 9. Videos were captured at 45 frames per second (fps) from the camera. We processed the video recordings with two different tracker configurations: one with anchor sets optimized empirically by an operator and one with automatic anchor set optimization. Both results are shown in Table 2.

**Figure 9:**
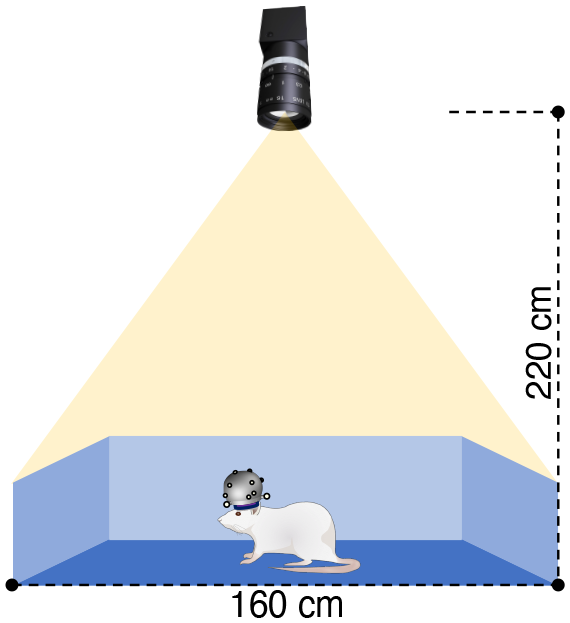
Reliability testing of the 3D tracking system was performed with a live rat subject. The visual target was mounted on the head on the animal. The rat was freely roaming in a 1.6 m by 1.6 m size arena that was placed under the camera at a distance of 2.2 m. In four recording sessions a total of ∼2.5 hours of data was recorded.

**Table 2:** Failure statistics of 3D tracking during reliability testing. The visual target is mounted on the head of a free roaming rat inside a 1.6 m by 1.6 m size rectangular arena. Failure rates are specified (in frame count and percentage of all frames) for both manually selected anchor sets (N = 9) and computationally optimized anchor sets (N = 9).

|  | Duration |  | Failure rate |  |  |  |
| --- | --- | --- | --- | --- | --- | --- |
|  |  |  | Manual |  | Optimized |  |
|  | [min] | [frm cnt] | [frm cnt] | [%] | [frm cnt] | [%] |
| Run 1 | 44.48 | 116,836 | 7,966 | 6.82 | 635 | 0.54 |
| Run 2 | 31.13 | 79,992 | 5,972 | 7.47 | 482 | 0.60 |
| Run 3 | 22.12 | 58,852 | 3,878 | 6.59 | 177 | 0.30 |
| Run 4 | 50.12 | 126,408 | 20,131 | 15.93 | 866 | 0.69 |
| Total | 147.85 | 382,088 | 37,947 | 9.93 | 2160 | 0.57 |

In the 2.5 hours long evaluation session, the tracker successfully tracked the 3D pose of the rat’s head on 99.4% of the video frames when automatic anchor set optimization was enabled. This compares to the 90.1% 3D tracking success rate when the anchor sets were manually optimized by an experienced operator. Fig. 10 shows the trajectory of the visual target mounted on the head of the animal during the 2.5 hour run time of the experiment with automatic anchor set optimization enabled. The spatial distribution of target locations where the tracker failed to provide 3D pose estimates appears to be sparse and uniform in the central part of the arena, but the density of failed detections is higher in or near corners. In the corners, the animals tended to rear up, groom, or otherwise occlude the view of the target from the camera, which resulted in fewer observations and valid anchor sets.

**Figure 10:**
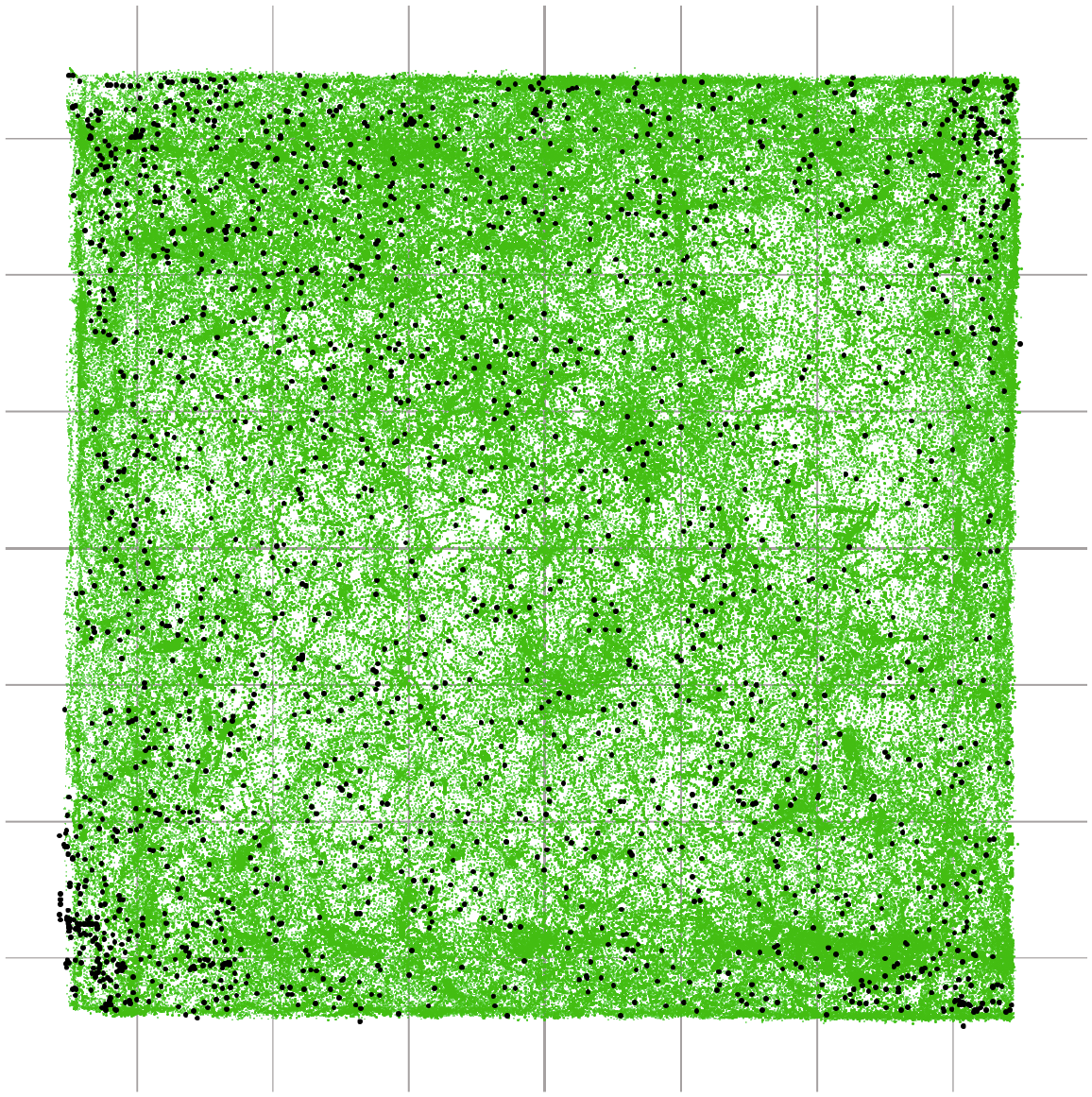
During reliability testing the rat was freely exploring a 1.6 m by 1.6 m size arena. The figure illustrates the trajectory of the rat’s head during the 2.5 hours of recordings. Small green dots represent the positions of the rat’s head on 379,928 video frames when the tracker succeeded in accurately calculating the 3D pose of the target. 2160 large black dots (size exaggerated for visibility) show the positions of the target when it was partially visible but the tracker was unable to determine its 3D pose. In these failure cases the tracker provides a position estimate based on the 2D position of the cluster of bright dots near the region of interest.

The videos were processed offline with simulated playback of the recordings at the original 45 fps. Average offline processing speed of the tracker was 44.4 fps with a minimum framerate of 10.0 fps. The lowest framerates are experienced immediately after a rapid change of orientation of the visual target that may force the tracker to evaluate a high number of anchor sets for correspondence matching. The evaluation was performed Apple MacBook Pro 16” equipped with a 2.3 GHz 8-core mobile Intel i9 CPU, running the Ubuntu 18.04 64-bit operating system on a virtual machine.

### 3.3. Comparison to ArUco Tracker

We compared the performance of a popular monocular 3D tracker solution to our system. ArUco is an ARTag-based tracker that is widely used in augmented reality, robotics, and scientific experiments. An ArUco marker is shown in Fig. 12. In order to provide a fair comparison to our tracker, we printed an ArUco marker of the same size, 54 mm by 54 mm, as the our visual target and tracked it with the same camera that we used for the evaluation of our tracking system.

**Figure 11:**
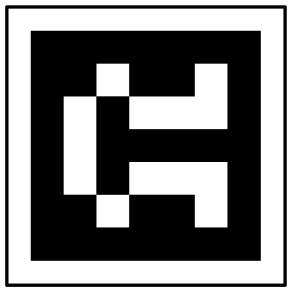
ArUco marker that was used to test the efficacy of the ArUco monocular optical tracker solution. The marker is printed on a sheet of paper and then fixed on a hard flat surface.

**Figure 12:**
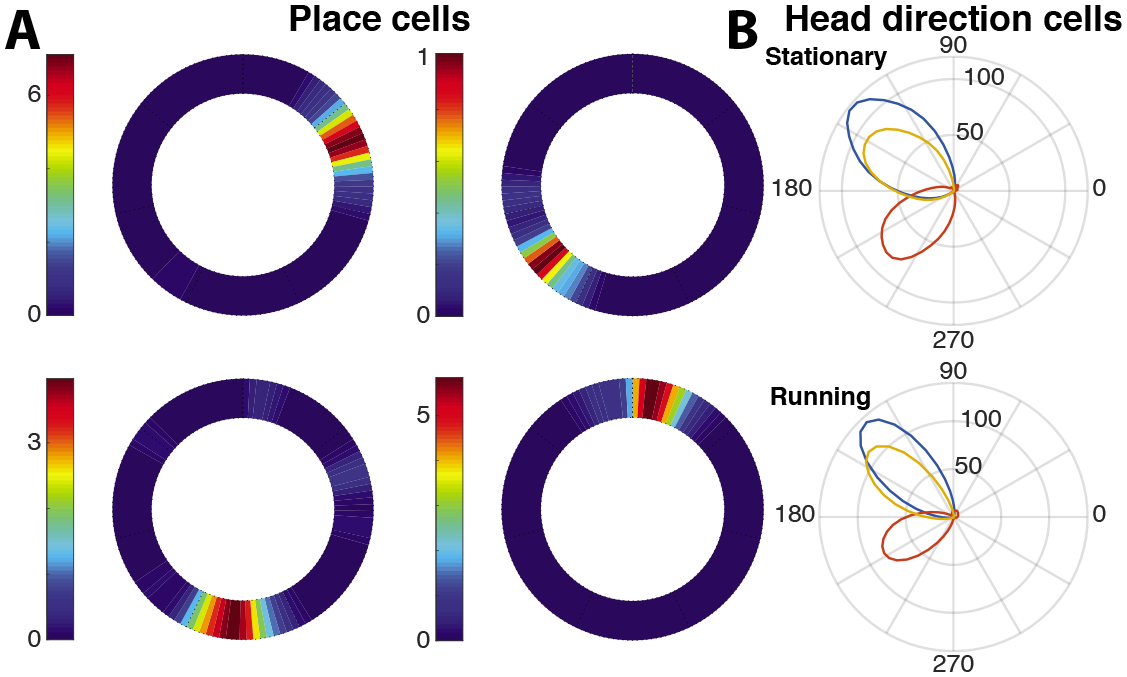
Application to place- and head-direction cell tuning. Examples of simultaneously recorded place and head direction cells from a single session of a rat running on a circular track. (A) Four example place cells. The annulus represents the circular track and color indicates occupancy-corrected firing rate in Hz (color bar indicates scale). (B) Polar plots of three head direction cells (red, blue, and yellow); radius represents occupancy corrected firing rate in Hz, and angle represents allocentric heading direction of the animal. The top plot shows the directional tuning when the animal is stationary and bottom plot shows tuning when running.

During testing, we managed to successfully track the marker with ArUco up to ∼1.5 m distance from the camera. When we moved the marker any farther, the tracker failed to provide any position or orientation estimate. Furthermore, even when the marker was closer than 1.5 m to the camera, ArUco only managed to locate it in an approximately ±70^◦^ range of angles relative to the front view. When the marker appeared at a sharper angle (*>* 70^◦^), the detection rate quickly plummeted and detection failed completely at around 75^◦^, which is consistent with results in other published literature [55].

Compared to ArUco, in our evaluations our proposed tracking solution had a success rate of 99.43% at 2.1 m distance from the camera and a range of supported orientations of up to ±120^◦^.

### 3.4. Application: Head Tracking during Hippocampal Recordings

To demonstrate the applicability of our tracking system in neurophysiological research, we performed chronic neural recordings from laboratory rats as they circumnavigated a circular environment. The neural recordings were performed using multi-tetrode hyperdrives [56]. While the complete set of recordings and experimental findings will be reported elsewhere, here we present examples that illustrate the applicability of the tracking system. **Fig. ??** shows the spatially selective neural activity of typical head direction cells and place cells collected using the tracking system. The sharp spatially selective tuning of place cells (with respect to position) and head direction cells (with respect to head orientation) provide demonstrative evidence of the utility of our pose estimation system for neurophysiological applications.

### 3.5. Application: Real-Time VR Manipulations

The application that initially motivated the development of our tracking system was its use in tracking the position and orientation of a rat while it circumnavigates a custom virtual reality (VR) Dome apparatus [51, 56]. The VR Dome—like many VR systems—requires an accurate estimate of the pose of the head, based on which manipulations of the visual scene can be performed. In our previously published experiments [56], the animal was harnessed to a radial boom arm that was connected to a centralized optical encoder to measure the angular position of the animal. On the basis of this, the visual scene could be adjusted. A major goal in developing the head tracking system was to overcome the need for the restraining harness, allowing free movement of the animal. However, the VR Dome imposes strict geometric constraints on camera-based tracking system. For example, the video images must be acquired from directly overhead, ∼1 m above the table on which the animal navigates. Moreover, the design of the Dome precludes the use of commercial systems that typically employ multiple cameras that record images from different perspectives (wide baseline). For details, see [51].

We successfully performed a set of experiments in 5 rats in which we changed the visual scene in response to the optically tracked position of the rat, while recording the activity of place cells. As discussed in [51], the switch to optical tracking reduced training time while maintaining the behavioral parameters observed during harnessed running. In addition, optical tracking provided us with the 3D orientation of the head of the animal, which are being analyzed for publication. The camera was capable of capturing 2048 × 2048 pixel frames at 90 fps, and the tracker was able to keep up with this frame rate up to 81 frames / sec (12 ms image processing pipeline). Ultimately, we chose 45 fps for the experiment as a balance between tracking reliability and data size. We tested the feedback latency of the apparatus (latency between movement of the rat and corresponding movement of the visual cues) with encoder-tracked and optical-tracked positions. The latency—including image capture, the entire tracking pipeline described above, and movement of the cues—was approximately 100 −110 ms for frame rates between 30 − 90 fps [51].

## 4. Discussion

We developed an open-source, monocular (single-camera) optical tracking system for animal research that consists of a small, compact, and lightweight visual target and software capable of tracking the target at a high frame rate in real time based on camera images. Our solution improves upon the state of the art by providing accurate 3D pose (position and orientation) at a wider range of orientations than other tracking systems. Performance evaluations using synthetic tests and laboratory rats demonstrated high accuracy and reliability. The use of a single camera and 3D printable visual target keeps setup simple and inexpensive, while maintaining flexibility for users to adapt the visual target for other applications with minimal effort.

In our extensive testing—benchmarking against a commercial system and several live animal experiments—our marker-based 3D tracking system proved to be more accurate, more robust, and effective over a larger range of angles than other state-of-the-art monocular tracking solutions. A significant advantage of the proposed solution is the extended range of trackable angles of the marker. Most other marker-based systems rely on markers that are only visible from a single side of the marker (visibility *<* 90^◦^), but our tracker is able to track its visual target in a significantly wider range of angles, up to ±120^◦^. This extended range enables the tracking of a larger variety of animal behavior than before.

The physical setup of the system is straightforward as it only requires a computer, single camera, a ring light, and the visual target that is mounted on the subject. The tracking software runs on Ubuntu Linux and uses the Robot Operating System (ROS) software framework to communicate with external applications. For data recording, playback, and visualization, ROS provides a set of convenient software tools that enable simple integration and debugging.

Accurate, real time, monocular pose estimation allows an experimenter to perform experiments where the specific pose of the animal can be used in closedloop to control behavioral parameters. As an example, these features are critical in a VR experiment where the rendered visual scene has to accurately reflect the quickly changing position and gaze direction of an animal, and the visual scene has to be generated and displayed with low latency and without jumps that would degrade the immersive experience. While markerless tracking methods exist, we believe the substantially improved reliability and accuracy of marker tracking is necessary in such sensitive applications. Being monocular and highresolution, our tracking algorithm can be implemented with minimal overhead in large-scale experiments that include other bulkier recording equipment. The algorithm can also be scaled to detect multiple targets with different marker configurations in the same camera frame, allowing for tracking of multiple features or social interactions.

Historically, neurophysiological recordings of cells such as place cells, head direction cells, grid cells, object cells, and other cell types in the rodent hippocampal formation have relied on video tracking systems. These systems typically calculate x-y position coordinates and head direction angles in the horizontal plane from the video image representing the orthographic projection of 2 or more sets of LEDs on the animals head onto the 2D camera sensor array. This technique has been sufficient for answering many of the basic questions about these cells, but increasingly sophisticated knowledge about the system (e.g., the 3D nature of head direction encoding [25]) and behavior require a finer-scale measurement of 3D pose. For example, animals exhibit a behavior called vicarious trial and error (VTE) at choice points on a maze, in which they move their heads back and forth in the directions of the two behavioral choices as they deliberate which way to proceed [57]. During VTE behavior, place cells transiently represent nonlocal trajectories down the different choice paths that correlate with the animals head direction. Traditional head tracking is sufficient to investigate this phenomenon on a 2D maze, but is not adequate to investigate behaviors in which the animals choices are between two paths that differ in the z axis (i.e., when the corresponding VTE behaviors are related to changes in head pitch rather than yaw). Another example of complex behavior is called head scanning, in which rodents pause and move their head back and forth to take in information about the external world during exploration [28]. Traditional, 2D head tracking techniques provide limited information to distinguish this type of head movement from the VTE behaviors displayed at choice points and other types of head movements that occur during behaviors such as grooming, object exploration, and rearing. Techniques such as those described here (perhaps in conjunction with other 3D sensors such as accelerometers) can provide a much richer trove of data that can be used to automatically characterize different aspects of complex behaviors, and thereby allow greater insight into the neurophysiological correlates of complex behavior and cognition.

## Acknowledgements

We thank F. Savelli, G. Secer, B. Krishnan, T. Danjo, S. Lashkari, and S. Leonard, for assistance with developing and testing the head tracking system.

## Contributions

**Balazs Vagvolgyi:** Conceptualization, Methodology, Software, Validation, Formal analysis, Investigation, Data Curation, Writing – Original Draft, Visualization. **Ravikrishnan Jayakumar:** Conceptualization, Methodology, Target design, Validation, Data collection and curation, Writing – contributions to draft, Visualization. **Manu Madhav:** Conceptualization, Methodology, Software, Validation, Formal analysis, Investigation, Data Curation, Writing – Original Draft, Visualization. **James Knierim:** Conceptualization, Writing – Reviewing and Editing, Supervision, Project administration, Funding acquisition. **Noah Cowan:** Conceptualization, Writing – Reviewing and Editing, Supervision, Project administration, Funding acquisition.

